# Lineage recording reveals dynamics of cerebral organoid regionalization

**DOI:** 10.1101/2020.06.19.162032

**Authors:** Zhisong He, Tobias Gerber, Ashley Maynard, Akanksha Jain, Rebecca Petri, Malgorzata Santel, Kevin Ly, Leila Sidow, Fátima Sanchís-Calleja, Stephan Riesenberg, J. Gray Camp, Barbara Treutlein

## Abstract

Diverse regions develop within cerebral organoids generated from human induced pluripotent stem cells (iPSCs), however it has been a challenge to understand the lineage dynamics associated with brain regionalization. Here we establish an inducible lineage recording system that couples reporter barcodes, inducible CRISPR/Cas9 scarring, and single-cell transcriptomics to analyze lineage relationships during cerebral organoid development. We infer fate-mapped whole organoid phylogenies over a scarring time course, and reconstruct progenitor-neuron lineage trees within microdissected cerebral organoid regions. We observe increased fate restriction over time, and find that iPSC clones used to initiate organoids tend to accumulate in distinct brain regions. We use lineage-coupled spatial transcriptomics to resolve lineage locations as well as confirm clonal enrichment in distinctly patterned brain regions. Using long term 4-D light sheet microscopy to temporally track nuclei in developing cerebral organoids, we link brain region clone enrichment to positions in the neuroectoderm, followed by local proliferation with limited migration during neuroepithelial formation. Our data sheds light on how lineages are established during brain organoid regionalization, and our techniques can be adapted in any iPSC-derived cell culture system to dissect lineage alterations during perturbation or in patient-specific models of disease.

## Main

Three-dimensional (3D) cerebral tissues derived from human induced pluripotent stem cells (iPSCs) – so-called cerebral organoids – mimic aspects of the in vivo architecture and multilineage differentiation observed in primary developing brain tissue^1^. Cerebral organoids also allow for the manipulation and analysis of human brain tissue in controlled culture conditions across time. Single-cell RNA-sequencing (scRNAseq) has been extensively used to unbiasedly identify molecularly distinct cell types in brain organoids^2–5^ as well as diverse other organoid systems. scRNAseq can also identify cells as intermediates between types, and cells can then be computationally aligned to delineate differentiation paths and order cells in pseudotime^6,7^. However, it is not possible to use these inferences to identify lineages.

Previous work in cerebral organoids has revealed the power of scRNAseq to understand brain region composition and progenitor-to-neuron differentiation trajectories in individual cerebral organoids. However, it has been difficult to understand how brain regions are established during organoid self-organization. Several lineage-coupled single-cell transcriptomics strategies have been employed to investigate clonal expansion and differentiation in mouse and zebrafish embryos as well as complex multicellular culture systems^8–12^. These efforts rely on either expression of reporter transcripts tagged with a unique sequence barcode^8,12^, or on CRISPR/Cas9 genomic scarring patterns generated by CRISPR/Cas9 genomic modification of expressed targets^9–11^. Lineage-coupled scRNAseq has allowed for better annotation of cell fate specifications and trajectory inferences in complex tissues and other cell differentiation scenarios.

Here, we have established a dynamic cell lineage recorder by coupling a highly complex barcode library together with an inducible Cas9 scarring system in iPSCs (Fig. 1a. From these iPSCs, we initiate organoids using approximately 2000 cells and at different stages of organoid development induce CRISPR-Cas9 genetic scarring to label progenitor cells that will give rise to distinct mature cell types with secondary lineage scars (Fig. 1b). We then use the barcodes and genetic scars together with the transcriptome profiles of single cells to reconstruct differentiation trajectories, trace cell lineages and explore the time points of fate decisions over the course of human cerebral organoid development from pluripotency, through neuroectoderm and neuroepithelial stages, followed by divergence into neuronal fates within forebrain, midbrain and hindbrain regions. This technique provides an opportunity to trace cell lineages in various 3D cell culture systems and opens up the possibility of dissecting lineage-specific alterations in patient-specific models of disease.

**Fig. 1.**
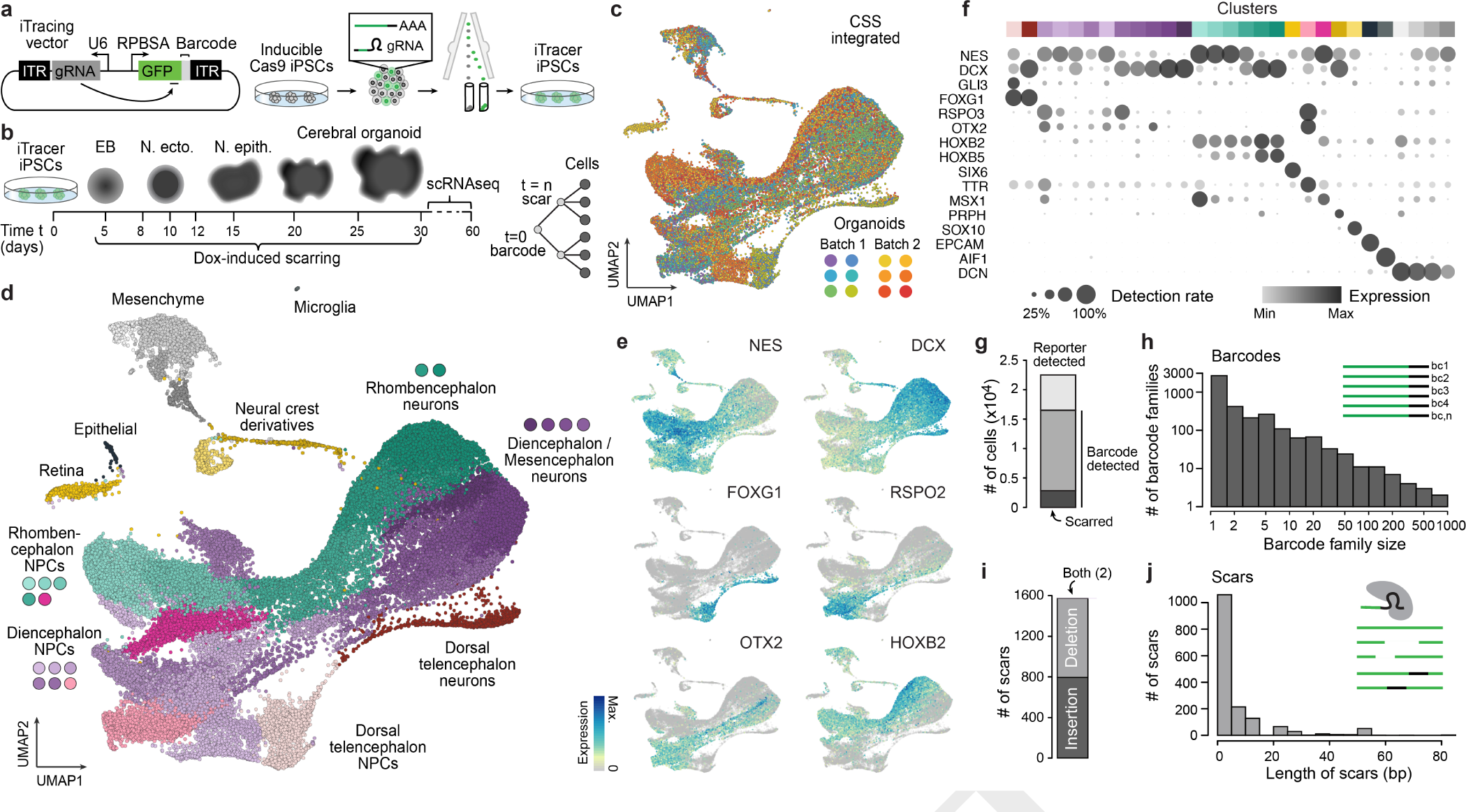
Single-cell transcriptome-coupled lineage recording in iPSC-derived human cerebral organoids using iTracer. a) Schematic of sleeping beauty vector used for lineage tracing. The reporter barcode is a random 10-mer. Experiments were performed using green and red fluorescent proteins, and the gRNA targets a location within the C-terminus of the fluorescent protein (see methods). The vector is introduced into iPSCs containing a doxycycline inducible Cas9 cassette and integrates through multisite undirected transposition. Fluorescent iPSCs are sorted and expanded prior to embryoid body (EB) formation. b) Scarring is induced at different time points through doxycycline induction of Cas9. c) Uniform Manifold Approximation and Projection (UMAP) embedding of scRNAseq data of 44,275 cells from 12 cerebral organoids after data integration using cluster similarity spectrum (CSS)16. Cells are colored by batch and organoid of origin. d) UMAP embedding with cells colored by cluster, and annotated with brain region or cell type identity. e-f) Expression of genes marking clusters observed in the organoids. g) Stacked barplot showing the number of cells where the iTracer reporter was detected (light grey), with only barcodes (dark grey) or barcodes and scars (black). h) Barplot of number of barcode families and sizes of families detected. i) Stacked barplot showing the number of scars created from insertions (light grey), deletions (dark grey) or both. j) Barplot of number of scars and scar lengths detected.

The lineage recording system, which we call iTracer, is based on the sleeping beauty transposon system which enables efficient transposition^13^ of exogenous DNA into multiple genomic loci within iPSCs. A poly-adenylated and barcoded (11 random bases) green or red fluorescent protein (GFP/RFP) reporter is driven by the RPBSA promoter. In the opposite direction 91 bases away from the RPBSA promoter, we have introduced a human U6 promoter driving a gRNA that targets a region within the 3’ portion of the GFP or RFP coding sequence. This construct is then introduced through electroporation into iPSCs that contain a doxycycline-inducible Cas9 cassette (iCRISPR)^14,15^. Fluorescent cells containing the reporter can be isolated using fluorescence activated cell sorting (FACS) before being propagated or cryopreserved for later use. Doxycycline introduction into the media induces Cas9 expression, followed by formation of Cas9-gRNA complexes leading to double-stranded break formation at the targeted location in the fluorescent reporter region of the recorder. These breaks are repaired by cellular machinery, which results in insertions and/or deletions, called scars, at the cut-site and can be read by sequencing the reporter transcript. Multiple barcodes and induced scars could in principle be detected per cell due to multiple insertions of the transposon-based reporter.

We visualized iTracer fluorescent reporter signals to confirm detection throughout organoid development (Extended Data Fig. 1a). Next, we used targeted amplicon sequencing of the barcode region to measure barcode diversity in a pool of iPSCs as well as from various stages of organoid culture. There was substantial barcode diversity in the iPSC pool, and the diversity remained relatively stable over the course of organoid development (Extended Data Fig. 1b). We used single-cell transcriptomes to determine the number of barcodes per iPSC and found an average of 2.85 barcodes per cell, with 81% of the measured cells having at least one barcode detected. The average group size of those cells that shared the same barcode was 1.54 cells (Extended Data Fig. 1c). Cells with a higher number of barcodes also had higher expression of the iTracer fluorescent reporter mRNA. These data indicated sufficient barcode complexity of iTracer originating iPSCs. We next analyzed the efficiency of Cas9-induced scarring at the embryoid body and neuroectoderm stages of cerebral organoid development by testing the duration and concentration of Doxycycline treatment (Extended Data Fig. 1d-e). We found scarring was most efficient when samples were incubated in 8 micrograms of Doxycycline for 24 hours and applied these conditions in all subsequent experiments. Together these data established suitable conditions for lineage-coupled single-cell transcriptomics in iPSC-derived cells and tissues using the iTracer system.

We next set out on a series of experiments using iTracer to study lineage dynamics during cerebral organoid development from pluripotency. Organoids were generated from approximately 2000 iPSCs containing the iTracer recorder. We induced scarring at day 4, 7, 15, 20 or 30, and performed scRNAseq after approximately 30, 45 or 60 days of culture (Fig. 1b). We used cluster similarity spectrum (CSS) analysis16 to integrate 44,275 cells from 12 organoids across two batches (Fig. 1c, Extended Data Fig. 2a-b, Supplementary Table 1 and 2), resulting in 29 cell clusters. Based on marker gene analysis and comparisons to primary reference atlases (see Methods), 90% of cells were annotated to central nervous system (CNS) cell types from dorsal telencephalon, diencephalon, mesencephalon, rhombencephalon, and retina, while a small fraction was annotated as non-brain populations including neural crest derivatives and mesenchymal cells (Fig. 1d-f, Extended Data Fig. 2c-f).

Overall, we detected iTracer readouts in 22,489 cells (51%) from this dataset (Extended Data Fig. 3a). Between organoids we detected a variable number of GFP/RFP reporter-expressing cells, barcodes, and scars (Extended Data Fig. 3a, Supplementary Table 1). Based on fluorescence, we noted that while the entire starting population of iTracer clones contained the reporter, reporter mRNA was detected in 51% of cells at the time of sequencing. Transgene silencing and sparsity of scRNAseq data are likely contributors to loss of the reporter detection^17,18^. Nonetheless, of those cells where the reporter transcript was detected by scRNAseq, we identified at least one barcode in 73% of cells (Fig. 1g, Extended Data Fig. 3a-b). Due to the nature of the sleeping beauty system, cells can have multiple iTracer insertions, allowing for unique barcode and scar compositions within individual cells. Within those cells where we detected at least one barcode, we found cells contained an average of 2.10 barcodes per cell. Groups of cells that share the same barcode composition, termed barcode families, ranged in size from 2 to 801 and averaged to approximately 11 family-members (Fig. 1h). CRISPR/Cas9 cuts resulted in a similar proportion of insertion and deletion scar types, which varied in overall length (Fig. 1i-j). Scarring efficiency also varied per organoid (Supplementary Table 1), yet of those cells where we detected a barcode, 17% were also scarred (Extended Data Fig. 3b). We found 237 groups of cells sharing the same scar and barcode composition, which were termed scar families.

We leveraged iTracer barcode and scar readouts, together with single-cell transcriptomes, to reconstruct cell type- and brain region-resolved lineage families in individual organoids (Fig. 2a-c, Extended Data Fig. 3c). We constructed organoid lineage plots, first traced by iPSC barcodes that were incorporated into each initializing embryoid body (EB), and second by the scars induced during the developmental time course. We analyzed the diversity of cell types within scar families where organoids were treated with Doxycycline at different time points. In one example we found cortical lineage cells clearly separated from others when the organoids were scarred at day 15 (Fig. 2d-e). Overall, we found a tendency where scarring at earlier time points lead to more diverse cell types within scar families than scarring at later time points (Fig. 2f). This increasing commitment through time reveals a coarse patterning window for brain regionalization, however sparse sampling of cells in whole organoids limits the depth of lineage analyses.

**Fig. 2.**
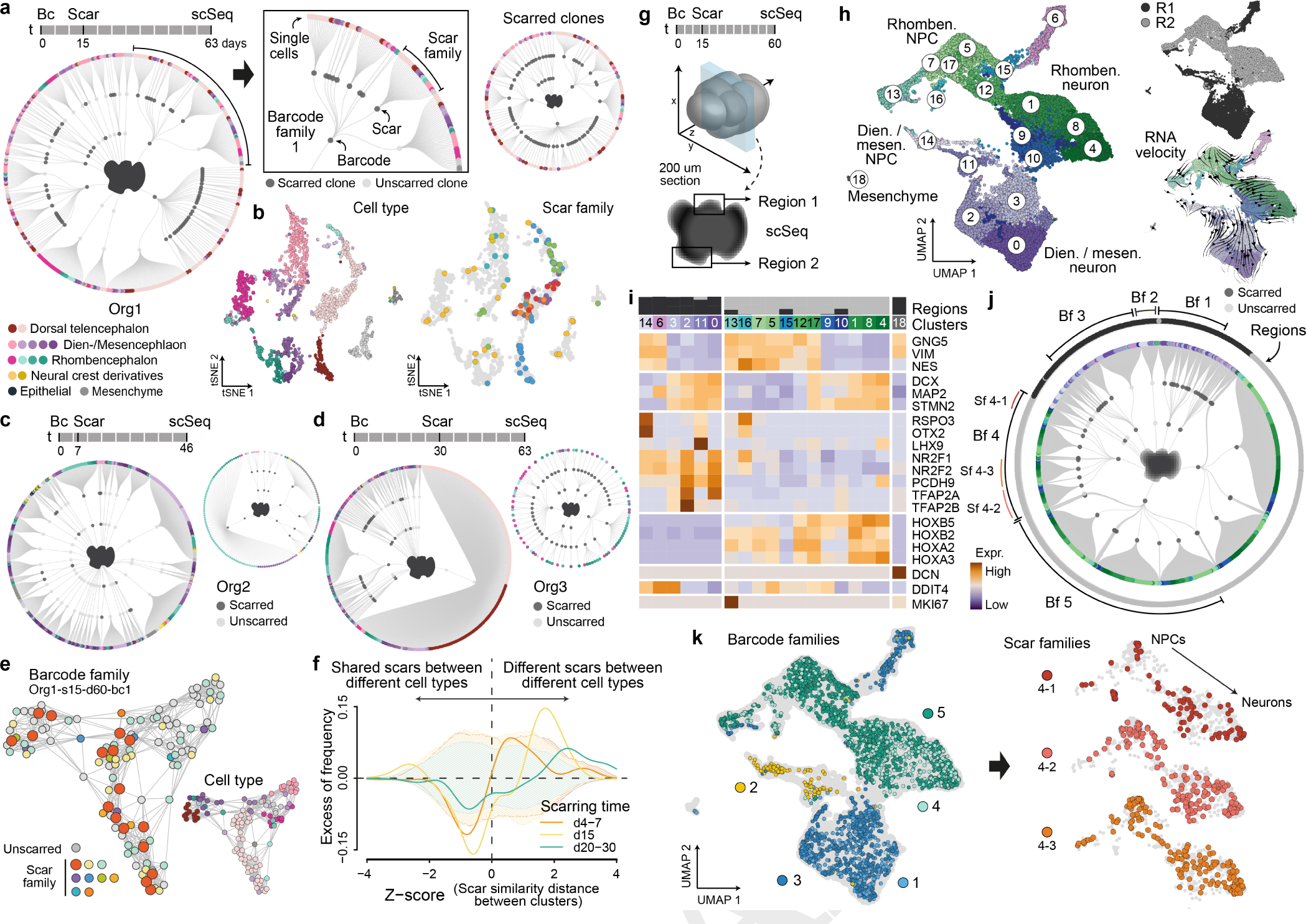
Organoid lineage reconstructions reveal regional clonality, a coarse patterning time window, and progenitor-neuron lineage families. a) Lineage plot shows full lineage reconstructions from a single organoid scarred at day 15 and sequenced at day 63 (left), as well as the subset of cells where scars were detected (right). The first and second order deviation nodes represent barcode and scar families, respectively, with the terminal branches indicating individual cells. Each cell is colored based on the cell type designation. b) tSNE embeddings show cells for this organoid colored by both cell type annotation and scar families (>=5 members). c-d) Lineage plots for two additional organoids scarred at day 7 and day 30, respectively. e) Scar pattern diversity in a single barcode family within Org1. Scar families in orange show enrichment in cortical brain regions. f) Frequency distribution of z-scores of scarring pattern distances between cell clusters, after subtraction of background distribution estimated by random sampling of scars. Different scarring times are shown separately. The dashed shadow backgrounds show the 90% confidence intervals, colored by the scarring time accordingly. This plot highlights that scarring at later time points separates lineages with different cell fates. g) Schematic of tissue section selection for deep-sampling. One 200um section was cut before selecting two spatially distant regions for microdissection. Single-cells were isolated from microdissected regions and processed for scRNAseq separately. h) UMAP embedding of scRNAseq data of 26894 cells from two microdissected regions within a single cerebral organoid scarred at day 15 and sequenced at day 60, cells are colored by cluster or originating region and annotated with brain regional or cell type identity. i) Expression heatmap showing brain region and cell type markers across clusters and regions shown in panel h. j) Lineage plot shows the lineage reconstruction combining both microdissected regions from the single organoid (left). The first and second order deviation nodes represent barcode and scar families respectively, with the terminal branches indicating individual cells. Originating region is annotated in the outer circle, with example barcode and scar families annotated. k) UMAP embedding of single-cells from the two microdissected regions colored by example barcode families indicated in panel j. Three different scar families from barcode family 4 are shown linking neural progenitor precursors to mature neurons.

Microdissection followed by scRNAseq allowed us to capture thousands of cells from distinct parts of the organoid thus increasing the sampling depth and the resolution of iTracer readouts. At the same time, it enabled us to understand the diversity of spatially distant regions at both the transcriptome and lineage level. To this end, we analyzed single-cell transcriptomes and iTracer readouts of two distinct regions that were microdissected from a 200 micrometer (um) tissue section originating from a 60-day-old organoid scarred at day 15 (Fig. 2g, Supplementary Tables 1 and 3). We found that these regions were transcriptionally divergent, where region one (R1) and region two (R2) were mainly composed of diencephalon/mesencephalon and rhombencephalon cell types, respectively (Fig. 2h-i, Extended Data Fig. 4a-b). We used iTracer readouts from these highly sampled regions to reconstruct barcode and scar families and identified entirely diverged cell lineages in R1 and R2 (Fig. 2j-k). We note that the iTracer lineage readouts enable connection of mature neurons to NPC pools (Fig. 2k); lineage connections that could not be resolved with transcriptome measurements or trajectory inferences alone (Fig. 2h). Furthermore, we found that transcriptome-based cell clusters formed three groups based on the enrichment of a distinct combination of barcodes in each group. Within two cluster groups (CG 1 and CG 2) we found a substructure based on enrichment of unique scar combinations and each sub-group represented a distinct cell state, indicating cell fate commitment before the time point of scarring (Extended Data Fig. 4d-g).

Intriguingly, we observed that within individual organoids, barcode families tended to accumulate in distinct brain regions. In particular, cells in CG 1 and CG 2, which each showed a distinct barcode composition, were mostly from different microdissected regions. To understand if this was a robust observation, we performed a permutation enrichment analysis using single cells from all lineage-traced organoids together to determine the likelihood that barcode families are shared between brain region-specific cell populations (Fig. 3a, Extended Data Fig. 5a-b). We identified three groups of cell clusters with significant enrichment of distinct barcode clones (Fig. 3b). Each group labeled distinct cell populations (Fig. 3c) indicating that indeed there is a robust trend during organoid development where initiating iPSC clones accumulate in distinct brain regional identities. Importantly, these results could not be solely explained by cell type composition differences in organoids (Extended Data Fig. 5c-d). We illustrated regional barcode accumulation by projecting four different barcode families on the overall UMAP embedding and saw that iPSC clones distribute into distinct brain regions or cell types (Fig. 3c-d).

**Fig. 3.**
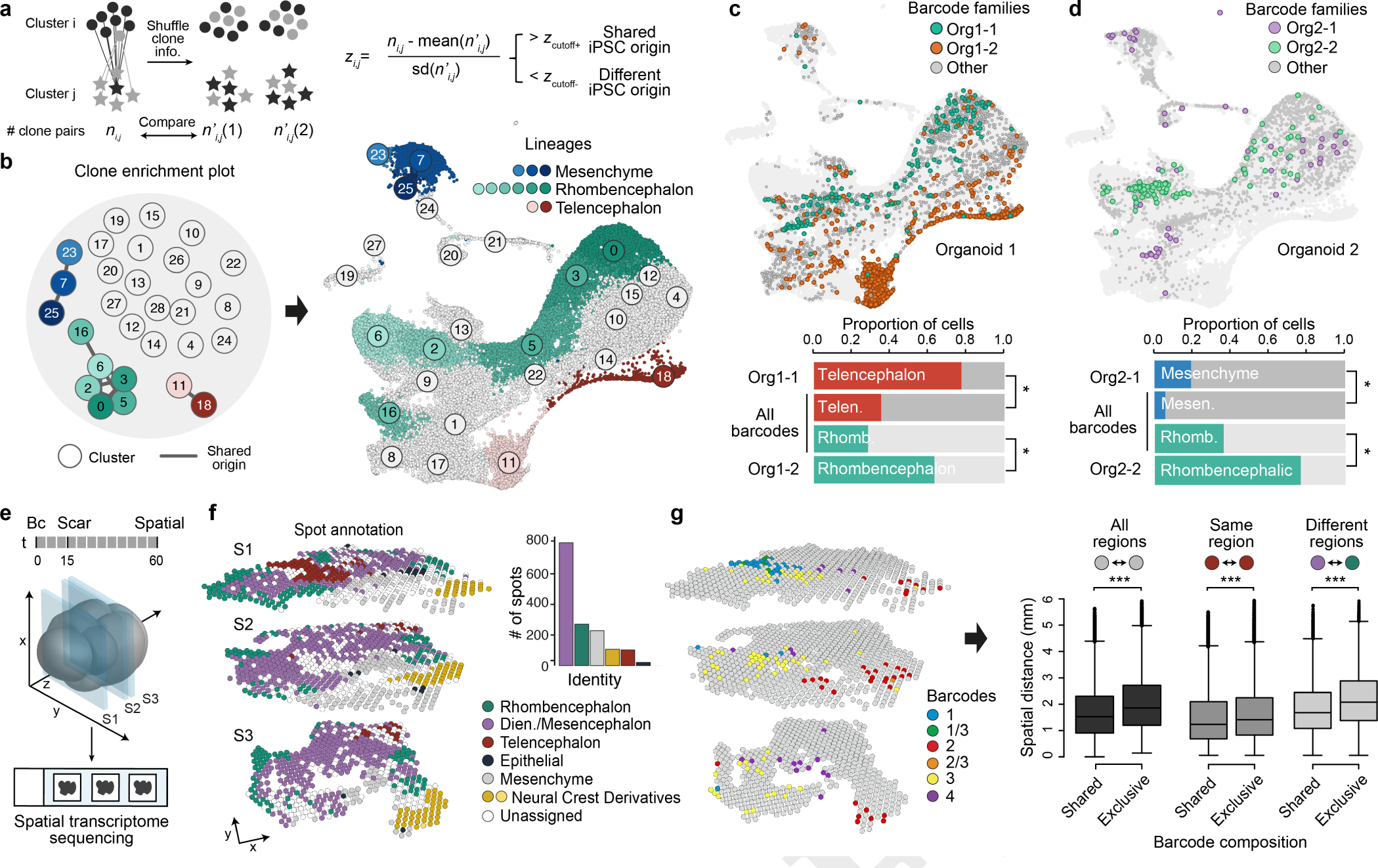
Spatial transcriptomics reveals that barcode families diversify and accumulate in distinct brain regions. a) Permutation enrichment analysis was used to determine the likelihood that barcode families would be shared between transcriptionally distinct clusters. b) Clone enrichment plot showing each cluster (circle), linked by edges representing significant enrichment of barcode clones, and colored by cluster identity (left). UMAP embedding of all whole-organoid scRNAseq data with cells labeled by cluster number and the cluster lineages colored where significant shared origin was detected. c-d) UMAP embedding is colored by 4 different clones, showing that iPSC clones tend to accumulate in distinct brain regions or cell types. Fisher’s exact test was performed comparing cell frequencies of a barcode family with all barcoded cells in the organoid in different clusters, * indicating p < 0.001. e) Schematic of tissue section selection for spatial transcriptomics. Three 10um frozen sections were adhered to a 10x Visum slide containing spots with barcoded capture probes before permeabilization of the tissue, cDNA synthesis, and subsequent construction of RNAseq libraries. f) 3D plot of spots corresponding to three capture tissue sections (S1-S3), colored by assigned cell type (CIBERSORTx). Barplot indicates the number of spots for each assigned cell type. g) 3D plot of spots across the three tissue sections where four example iTracer barcodes are highlighted, indicating lineage regionalization. Boxplots of the iTracer barcode composition of each spot vs. the spatial distance of each spot found in the three sections. Regions that are spatially close to each other share similar barcode composition, whereas barcode similarity decreases as spots increase in spatial distance. This occurs when comparing any two spots regardless of spot annotated brain region (left), between any two spots within the same annotated brain region (middle), and between any two spots which are assigned to different annotated brain regions (right). Two-sided Wilcoxon rank sum tests were performed comparing shared to same and exclusive groups, *** indicates p-values < 0.0001.

We hypothesized that the enrichment of barcodes in distinct brain regions may be due to the spatial arrangement of cells throughout the organoid. To understand the spatial distribution of barcodes within organoids we used spatial transcriptome sequencing based on the 10x Visium platform to measure gene expression and iTracer readouts from three intact 10um tissue sections within a 62-day-old cerebral organoid that was scarred at day 15 (Fig. 3e, Supplementary Table 1 and 4). Briefly, each tissue section was adhered to a slide, specifically to a 6.5 × 6.5-millimeter capture area containing several thousand spots, where each spot contains millions of capture oligonucleotides each with a unique spatial barcode. Tissue sections were first permeabilized to allow for mRNA capture, before captured sequences were amplified and sequenced. We obtained transcriptome, barcode, and scar topology of 2038 spots, where each spot can contain up to 10 cells. We used a digital cytometry approach (CIBERSORTx)^19^ to deconvolute and assign each spot to cell populations identified within cerebral organoids using scRNAseq (Fig. 3f, Extended Data Fig. 6a-g). We identified spots assigned to telencephalon, diencephalon/mesencephalon, and rhombencephalon, as well as non-cerebral identities. Spots with the same annotation were generally clustered together within the same slice and persistent throughout the depth of the organoid, revealing transcriptomic regionality. In total 41.7% of spots recovered contained at least one barcode, consistent with what we observed with scRNAseq barcode capture (Supplementary Table 1). Spots with the highest barcode detection were also co-localized with the highest expression of the iTracer RFP reporter (Extended Data Fig. 6h-i). We observed lineage regionality as iTracer barcodes exhibited distinct spatial distributions within the same slice (Fig. 3g). Barcodes detected within spots tended to overlap less with increasing physical distance between spots and this pattern was consistent irrespective of cell type assignment (Fig. 3g, Extended Data Fig. 6j-k). Together, lineage-coupled single-cell and spatial transcriptomics revealed that organoid brain regions have distinct lineage compositions that can be traced back to clonality within the initializing embryoid body.

The iTracer system provided comprehensive information of multiple static snapshots of the lineage formation process. However, this system lacks the dynamic information required to understand the spatial restriction of lineages and their expansion in a developing organoid. We speculated that the observed distinct lineage composition of brain regions could occur due to early spatial restrictions of clones at the EB stage and subsequent local amplification. Therefore, as an independent and complementary method that can provide information on cell lineage dynamics, we performed long term live imaging of developing cerebral organoids using four-dimensional (4D) lightsheet microscopy in order to directly track lineages over time (Fig. 4a, Extended Data Fig. 7a, Supplementary Movie 1-2). Briefly, we generated organoids containing 5% iPSCs that have nuclei labelled with a uniform fluorescent reporter, FUS-mEGFP^20^, and imaged the sparsely labeled organoids with an inverted lightsheet microscope (Fig. 4a). EBs were embedded in matrigel in the imaging chamber, cultured in neural induction media and the development was tracked for 65-100 hours (Fig. 4b, Supplementary Movie 1). As the EB grows and develops, we observed the formation of seal lumens, each of which can be tracked in 3 dimensions (Fig. 4c). We tracked the lineage of a single nucleus throughout the recording time using a new large-scale tracking and track-editing framework Mastodon, a Fiji^21^ plugin, that allows for semi-automated tracking and curation of nuclei lineages in large 4-Dimensional (4D) datasets (Fig. 4d). We visualized the spatial distribution of daughter cells derived from the originating nucleus, which we call lineage one (L1), and generated a lineage tree resulting from 100 hours of proliferation (Fig. 4f, Extended Data Fig. 7b, Supplementary Movie 3). We observed that L1 remained confined to the same lumen area throughout the recording time (Fig. 4d, Extended Data Fig. 7c). We tracked three additional nuclei, where two nuclei were neighbors within the same lumen area as L1 (L2-3) and the third nucleus (L4) was positioned diametrically opposite in a distinct future lumen area (Fig. 4g, Extended Data Fig. 7c-d, Supplementary Movie 4). We quantified the spatial distances between each tree, and inspected the distribution of all the daughter cells within the organoid three-dimensional (3D) space (Fig. 4g-h). During the course of 65 hours, originating nuclei gave rise to 13 daughter nuclei on average, which all populated the expanding organoid but remained spatially restricted to the parent lumen and exhibited limited migration away from their lineage members (Fig. 4g-h, Extended Data Fig. 7c-f). These results assert a link between our previously observed brain region clone enrichment and the position in the neuroectoderm (Fig. 4i). Altogether, this shows that cell lineages develop with restricted spatial positioning in the organoids followed by brain patterning and cell fate commitment.

**Fig. 4.**
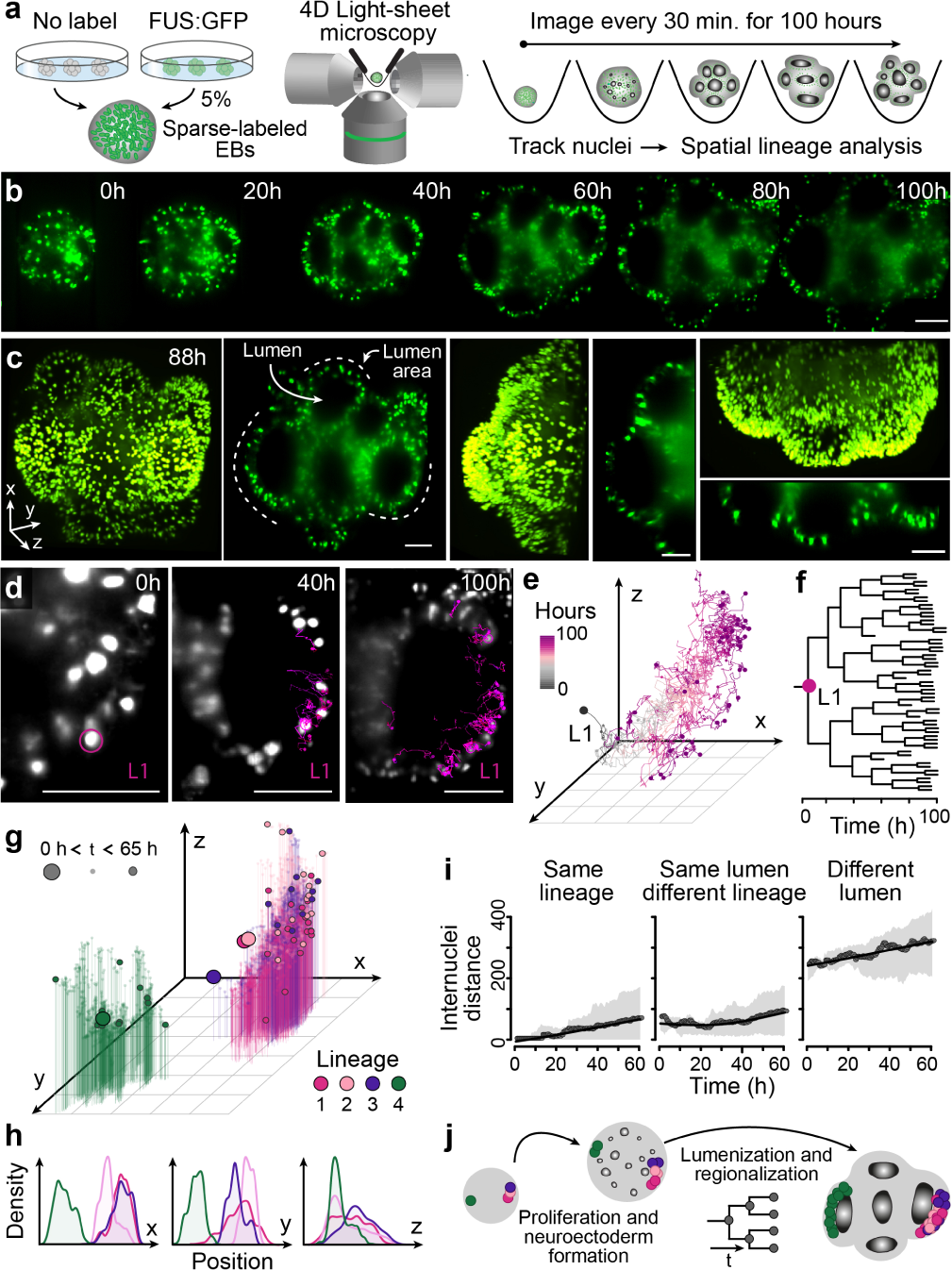
Long term nuclei tracking in cerebral organoids using light-sheet microscopy enables the study of spatial lineage dynamics. a) Schematic showing the experimental setup. Organoids were initiated by aggregating unlabeled (WTC) and fluorescently-tagged (FUS-mEGFP) iPSC lines to generate sparsely-labeled embryoid bodies. Developing organoids were imaged at 18.5x magnification over a 100-hour time course starting at 4 days post aggregation. The cerebral organoid protocol was slightly modified in order to accommodate imaging (see Methods). Scale bar is 100 μm in all images. b) Cross sections (x-y) still images from 0-100-hour time points. Labeled nuclei are colored in green. c) 3D projection and cross-sections of the x-y, x-z and y-z plane at 88 hours. Empty lumen cavity and lumen areas are annotated on the image. d) Selected images show the increment in nuclei tracks of lineage one (L1) from three timepoints over 100 hours. e) 3D plot showing spatial distribution of all nuclei in L1 over 100 hours. f) Lineage tree across time for L1. g) 3D scatterplot of spatial distribution of nuclei from four lineages (L1-L4) traced over 65 hours. Big, medium, and small dots represent time point 0 hours, 65 hours, and times between 0 and 65 hours, respectively. h) Density plot showing the distribution of all four lineages in x-y-z planes. i) Scatter plot showing the internuclear distance between any two nuclei in the same lineage (L1/L2/L3/L4), different lineages in the same lumen (L1-L3) and for nuclei in different lineages (L1/L2/L3 and L4). j) Illustration shows the proliferation and regionalization of tracked lineages in the organoid over 100 hours.

Here we present an extensive examination of lineage dynamics in the developing cerebral organoid. Using iTracer, we were able to successfully reconstruct sparse lineages within whole organoids at various stages of development. We observed a coarse timing for cell fate commitment and a robust pattern of clonal accumulation in distant brain regions. Transgene silencing, single-cell dropout, and depth of sampling proved to be hurdles in collecting complete lineage information from the iTracer system. Despite these limitations we show that by increasing sampling we can elucidate lineage connections between specific NPCs and mature cell types, as well as reveal timing of cell state emergence with scars. We resolve gene expression and lineage regionality in cerebral organoids using iTracer-coupled spatial transcriptomics. Furthermore, the data generated using 4D light sheet microscopy suggests that brain regionality observed in the two-month organoid is linked to the position in the neuroectoderm, followed by local proliferation with limited migration during neuroepithelial formation. Regionalization patterns then emerge and spatial constraints lead to the observed clonality in regionalized and differentiated cell fates. In the future, lineage tracking using iTracer and long term 4D light sheet imaging will be powerful methodological approaches to understand the effect of mutations that disrupt brain regionalization and other neurodevelopmental disorders.

## AUTHOR CONTRIBUTIONS

T.G., B.T., J.G.C. designed the iTracer system. R.P., T.G., L.S., S.R. established the iTracer system. M.S., R.P., A.M. performed FACS. R.P., A.M., A.J., F.S.C. performed organoid culturing. R.P., A.M., T.G. performed bulk and single-cell genomic experiments. A.M., J.G.C. performed organoid microdissection experiments. A.M. performed organoid spatial transcriptomics experiments. Z.H., T.G., A.M. analyzed the sequencing data. A.J. performed the lightsheet microscopy. A.J., K.L., M.S., Z.H. performed the nuclei tracking and data analysis. Z.H., T.G., A.M., A.J., J.G.C, B.T. designed the study and wrote the manuscript.

## DATA AVAILABILITY

Sequence data that support the findings of this study have been deposited in ArrayExpress with the accession codes E-MTAB-xxxx (bulk RNA-seq of barcode libraries), E-MTAB-xxxx (single-cell RNA-seq data based on 10x Genomics) and E-MTAB-xxxx (spatial transcriptomics RNA-seq data based on 10x Visium). Processed sequencing data have been deposited in Mendeley Data with doi http://doi.org/10.17632/nj3p3pxv6p.1.

## CODE AVAILABILITY

The computational code used in this study is available at GitHub 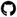(https://github.com/quadbiolab/iTracer) or upon request.

## ACKNOWLEDGEMENTS

We thank Jan Philipp Junker, Bastiaan Spanjaard and Bo Hu for inspiring discussions and protocols. We would also like to thank Andrea Boni and Petr Strnad for helping us in acquiring movies of developing organoids with their lightsheet system. Additionally, we would like to thank Pavel Tomancak, Vladimir Ulman, and Tobias Pietzsch for their support in implementing the Mastodon plugin for tracking Lightsheet data. Illumina sequencing was done in the Genomics Facility at D-BSSE, ETH Zurich and by Barbara Schellbach and Antje Weihmann at the Max-Planck-Institute for Evolutionary Anthropology. FACS sorting support was provided by the single-cell facility at D-BSSE, ETH Zurich. This project has been made possible in part by the Chan Zuckerberg Initiative DAF (grant CZF2017-173814, JGC and BT), an advised fund of Silicon Valley Community Foundation, the European Research Council (Anthropoid-803441, JGC; Organomics-758877, BT; Braintime-874606, BT), the Swiss National Science Foundation (Project Grant-310030_84795, JGC) and the National Center of Competence in Research, Molecular Systems Engineering.

## Materials and Methods

### Establishment of dynamic cell lineage reporter vector

iTracer plasmids were constructed by modifying sleeping beauty reporter plasmids pSBbi-GH and pSBbi-RH13. pSBbi plasmids were a gift from Eric Kowarz (Addgene plasmid 60514/ 60516; http://n2t.net/addgene:60514; http://n2t.net/addgene:60516; RRID:Addgene_60514; RRID:Addgene_60516). Plasmid generation consisted of 2 steps with initially removing the hygromycin cassette and inserting a 11 bp barcode tag in the 3’ UTR region of the fluorescent reporter gene. Secondly, the gene of interest (GOI) site including its promoter region was replaced by a human U6 promoter driving the expression of a gRNA that targets the fluorescence gene. In detail, a long-range PCR was performed to amplify and extract the plasmid backbone excluding the hygromycin cassette from pSBbi plasmids following manufacturer recommendations (Phusion Hifi Ready Mix 1x, 50 ng pSBbi, 0.2 uM of each primer (primers provided in Supplementary Table 5), DMSO 3%, 50 uL total, 25 cycles). In order to remove original plasmid from the backbone, the PCR reaction was digested by a combination of restriction enzymes directly added to PCR reaction (37uL H2O, 10 uL ThermoFisher FD Buffer, 1uL each FD enzyme DpnI, CpoI, Esp3I, 37C 1h). Reactions were purified using QIAGEN PCR Purification kit. The purified backbone was then used for performing a Gibson assembly following the instructions described in the Crop-Seq manual22 in order to introduce barcodes by using an oligo containing random nucleotides (Supplementary Table 5). The manual was only adapted such that bacteria were not plated after recovery but were instead completely transferred to an overnight-culture with 1x Ampicillin to maintain barcode heterogeneity. Plasmid DNA was isolated using the QIAGEN MiniPrep Kit. CropSeq plasmids containing a gRNA targeting either dTomato (GGTGTCCACGTAGTAGTAGC) or GFP (TGTTCTGCTGGTAGTGGT) were generated following protocol instructions (oligos provided in Supplementary Table 5). These plasmids acted as templates to amplify and extract the human U6 promoter, the gRNA and the gRNA scaffold region using the PCR conditions as described above. Primers (Supplementary Table 5) compatible with subsequent cloning were used and the purified PCR product was digested with ThermoFisher FD BshTI and FD PaeI prior to ligation. The backbone for cloning was obtained by digesting plasmids containing barcodes with ThermoFisher FD BshTI and FD PaeI. To guarantee that there is no ligation of the cut region back to the plasmid, the backbone digest was run on an Agarose gel and the backbone region was excised and purified. Backbone and insert were ligated using NEB QuickLigase and E. coli was subsequently electroporated with the ligation product following the CropSeq manual. Recovered cells were again directly transferred to an overnight culture and the plasmid DNA was extracted and purified using QIAGEN MediPrep.

### iCRISPR cell line and assessment of scarring efficiency

We used the iCRISPR iPSC cell lines with inducible Cas9 created previously as described14. DNA from iCRISPR was sent to Cell Guidance Systems Genetics Service Cytogenetics Laboratory and tested for copy number changes using the Agilent ISCA 8×60K v2 array. Array analysis revealed three apparently clonal DNA copy number changes. Gain of the entire long arm of chromosome 1 was detected and estimated to be present in about 70% of the cells. An approximately 3.7Mb DNA copy number loss of the proximal short arm of chromosome 19, band p12, was detected and present in about 30% of the cells. A commonly found mutational gain of approximately one megabase was detected within the proximal long arm of chromosome 20, band q11.21. Finally, inferCNV (https://github.com/broadinstitute/inferCNV) on the scRNAseq data of iCRISPR suggested a gain of a large portion of chromosome 12 in a subset of the cells.

To test the scarring efficiency upon Doxycycline induction, iCRISPR cells were cultivated using standard feeder-free conditions in mTeSR1 (StemCell Technologies) on matrigel-coated plates. Cells were nucleofected with 10ug lineage recorder DNA and 1ug Sleeping Beauty transposase following the manufacturer’s protocol and using the B-16 program of the Nucleofector 2b (Lonza) in cuvettes for 100μl Human Stem Cell nucleofection buffer (Lonza, VVPH-5022). Nucleofection reactions were plated on matrigel-coated plates and allowed to expand for 5-7 days before FACS. To select for cells with successful integration of the cell lineage reporter, RFP/GFP+ cells were sorted into 1.5mL tubes (~120,000 cells total) and plated in 12-cell matrigel-coated plates with mTeSR1 and Rock inhibitor (1:250). Following cell recovery, we used 2ug of Doxycycline15 for 0, 1, or 2 days before changing back to mTeSR1 base media. Optimization of Doxycycline incubation on 3D-cultures was done at EB and Neuroectoderm stages across 0.5, 1, 2, 4, and 8ug concentrations of Doxycycline for 24-hour incubations, before changing back to mTeSR1 and NIM base medias, respectively. iPSCs were harvested using Accutase (Sigma Aldrich) for 5-7 minutes before quenching with Knock-Out Media (ThermoFisher Scientific) and centrifuging at 200g for 5 min. 100,000 iPSCs were taken for DNA extraction using Quick Extract (Lucigen), whereas organoid samples were directly added to Quick Extract (Lucigen), and vortexed. All samples were vortexed shortly before heating to 65°C for six minutes, before vortexing again and heating to 98°C for two minutes. We used 50ng of input DNA for scar region amplification (primers provided in Supplementary Table 5). Quality of amplified product was checked with a 2% E-Gel (ThermoFisher Scientific) before Illumina sequencing adapters were added in a subsequent PCR reaction. Bulk scar libraries were cleaned with magnetic beads (Beckman Coulter) before checking quality with a 2% E-Gel. Libraries were sequenced on Illumina MiSeq Nano. Scar detection was performed using CRISPRESSO^23^.

### Preparation of Organoids and Scarring

iTracer+ iCRISPR cells were prepared as previously described above. Following cell recovery after FACS, 2000 cells per well in a 96-well plate were seeded and differentiated into cerebral organoids using a whole organoid differentiation protocol1,4. Throughout development organoids were scarred by activation of inducible Cas9 (Fig. 1b, Supplementary Table 1). Scarring was achieved by first selecting the organoid to be scarred and transferring it to a 6-24-well plate (depending on organoid size) filled with 8ug Doxycycline scarring media (base media depending on age of organoid, Supplementary Table 1). Organoids were incubated in scarring media for 24 hours before returning to base media without Doxycycline.

### Bulk barcode detection

IPSCs and 19 organoids ranging in stage from EB to day 30 were used for bulk analysis to assess the capture and diversity of iTracer barcodes. We propagated and harvested samples similarly to how we described above. Briefly, iCRISPR cells were cultivated using standard feeder-free conditions in mTeSR1 (StemCell Technologies) on matrigel-coated plates. Cells were nucleofected with 10ug lineage recorder DNA and 1ug Sleeping Beauty transposase following the manufacturer’s protocol and using the H9 program of the 4D-Nucleofector (Lonza) in cuvettes for 100μl Human Stem Cell nucleofection buffer (Lonza, VVPH-5022). Nucleofection reactions were plated on matrigel-coated plates and allowed to expand for 5-7 days before FACS. To select for cells with successful integration of the cell lineage reporter, RFP/GFP+ cells were sorted into 1.5mL tubes (120,000 cells total) and plated in 12-cell matrigel-coated plates with mTeSR1, Rock inhibitor (1:250), and Primocin (1:250). DNA isolation and bulk barcode libraries were prepared as we did bulk scar libraries (primers provided in Supplementary Table 5). Libraries were sequenced on Illumina MiSeq Nano. Analysis was done using a custom perl script to count the frequency of each uniquely detected barcode.

### Preparation of single-cell transcriptomes from whole lineage traced organoids

Whole organoids were dissociated for generating single-cell gene-expression libraries. In brief, organoids were transferred to HBSS (without Ca2+ and Mg2+, −/−) and cut into two pieces to clear away debris from the centre of the organoid (2–3 washes in total). Organoid pieces were then dissociated using Neural dissociation kit (P) using Papain-based dissociation (Miltenyi Biotec). Organoid pieces were incubated in Papain at 37°C (enzyme mix 1) for an initial 15 min, followed by addition of Enzyme A (enzyme mix 2) to the Papain mix. Organoid pieces were then triturated using wide-bore 1,000-ml tips and incubated for additional intervals of 5–10 min with triturations between the incubation steps, amounting to a total Papain incubation time of approximately 45 min. Cells were filtered through a 30-μm strainer and washed, centrifuged for 5 min at 300g and washed 3 times with HBSS (−/−). Cells were filtered through a 20μm strainer and washed, centrifuged for 5 min at 300g and washed 3 times with HBSS (−/−). Resulting cells were then assessed (count and viability) using Trypan Blue assay, counted using the automated cell counter Countess (Thermo Fisher). Finally cells were diluted to an appropriate concentration to obtain approximately 5,000-7,000 cells per lane of a 10x microfluidic chip device. Single-cell cDNA was synthesized per manufacturer recommendations before continuing to library preparation with 25% of the total cDNA volume. Libraries were sequenced on Illumina NovaSeq S1 and on Illumina HiSeq 2500.

### Barcode and scar detection from single-cell cDNA

Barcode and scar regions were amplified from 60-70ng of cDNA remaining from the single-cell RNAseq preparation with three separate PCR reactions. First cDNA was amplified via PCR broadly targeting a region containing both the scar and barcode. Subsequently, the reaction was split equally and we performed a nested PCR separately targeting the barcode and scar regions (primer sequences are provided in Supplementary Table 5). Lastly, we added Illumina sequencing adapters. Following every PCR reaction the samples were cleaned-up using magnetic beads (Beckman Coulter) and libraries were sequenced on Illumina NovaSeq S1.

### Alignment of single-cell transcriptomes and iTracer readouts

We used Cell Ranger (10x Genomics) to demultiplex base call files to FASTQ files and align reads. Default alignment parameters were used to align reads to a modified human reference including the fluorescent reporter (GFP/RFP) from the cell lineage recorder (hg38). Barcode and scar libraries generated from 10x cDNA were also aligned using default parameters with the exception that the force-cells argument was set to 200,000 and were aligned to a custom reference of the region of interest. This reference was constructed following 10x recommendations.

### iTracer readout filtering

iTracer barcode transcripts are first filtered at the UMI level, where transcripts are only retained should they have more than three reads. We then plotted the distribution of reads per UMI against the frequency of read depth per UMI and fit a line with loess through that distribution (*loess*(*nReads log*(*Freq_n_Reads*))) where values smaller than one were set to one. We then calculate the first minimum and removed everything which had smaller coverage than this point. iTracer barcodes began and ended with As or Ts, barcodes that did not match this pattern were removed. We filtered barcode transcripts so that when we detected the same UMI for different barcodes in the same cell the one with the highest read coverage was retained. Furthermore, when transcripts with the same UMI and barcode were found in multiple cells the transcripts were removed from these cells. Barcodes were further filtered so that when any barcodes with a hamming distance of one within any single cell were found, the barcode with the highest coverage was retained. Lastly, if more than 10 barcodes were detected in the same cell, the cell was ignored for lineage reconstructions.

iTracer scar transcripts are filtered similarly, where transcripts were first filtered at the UMI level, again we only retained those transcripts that had more than three reads. As we did for the barcodes, we plotted the distribution of reads per UMI against the frequency of read depth per UMI and fit a line with loess through that distribution where values smaller than one were set to one. We calculated the first minimum and removed everything which had smaller coverage than this point. We also filtered transcripts so when we detected the same UMI for different scars in the same cell, the one with the highest read coverage was retained. Transcripts were further filtered out if they had the same UMI and scar but were found in multiple cells. Lastly, we only kept scar transcripts where the same UMI was found in barcode and scar libraries. Similarly, barcode transcripts without corresponding scar transcripts were also excluded.

### Analysis of whole-organoid single-cell RNA-seq data

Seurat (v3.1)^24^ was applied to the scRNAseq data for preprocessing. Ribosomal protein genes and pseudogenes were excluded from the analysis. Generally, cells with more than 6,000 or less than 600 detected genes, as well as those with mitochondrial transcript proportion higher than 20% were excluded (Supplementary Table 1). After log-normalization, 5,000 highly variable genes were identified using the default vst method, where cell cycle related genes were excluded (Supplementary Table 6). Cell cycle scores were then calculated and regressed out from the highly variable gene expressions to reduce its confounding effect. The regressed-out expression levels were then z-transformed, followed by principal component analysis (PCA) for dimension reduction. Uniform Manifold Approximation and Projection (UMAP) was applied to the top-20 principal components (PCs) for visualization.

To integrate data of different organoids, Cluster Similarity Spectrum (CSS)16 was calculated as described. In brief, cells from each organoid were subset, and louvain clustering (with resolution 0.6), implemented in Seurat, was applied based on the pre-calculated top-20 PCs. Average expression of the pre-defined highly variable genes was calculated for each cluster in each organoid. Afterwards, Spearman correlation coefficient was calculated between every cell and every cluster in all organoids. For each cell, its correlations with different clusters of each organoid were z-transformed. Its z-transformed similarities to clusters of different organoids were then concatenated as the final CSS representation. UMAP and louvain clustering (with resolution 1) was applied to the CSS representation. Cluster annotation was done by combining expression patterns of canonical cell type markers, e.g. NES, DCX, SIX6, AIF1, DCN and EPCAM, and VoxHunt^4,25^ to compare the average transcriptome of clusters to different mouse brain regions.

For each organoid, a four-layer lineage tree was reconstructed. The pseudo-root node, representing the organoid, was considered as the first layer. Barcode families, i.e. cells with the same barcode combination detected, were considered as the second layer. Cells in the same barcode family were likely expanded from the same iPSC. In each barcode family, scar families, i.e. cells with the same scar combination, were considered as the third layer, which represent cells in the organoid expanded from the same cell when Cas9 was induced. At the end, cells were considered as the fourth layer. The lineage trees were visualized via the radialNetwork function in the networkd3 R package.

### Quantification of scar family composition differences between clusters

To quantify scar family composition differences between clusters, proportions of cells in different scar families of each barcode family were calculated. To increase the robustness of the estimate, one pseudocount was added to each scar family before calculating the proportions. Here, only barcode families satisfying the following criterias were considered: 1) contain at least five scarred cells; and 2) the second most frequent scar contains at least 10% of scarred cells in the barcode family. For each cell cluster, its scar family proportions of different barcode families in different organoids were concatenated to represent its scar family composition (denoted as *s_i_*). The scar family composition differences between two clusters *i* and *j* (denoted as *d_i,j_*) was then defined as the Euclidean distance between *s_i_* and *s_j_*.

In order to estimate the statistical significance of the scar family distance between two clusters, 1,000 random shuffling of scars were done. During each shuffling, the scar information of cells in the same barcode family were randomly shuffled. Afterwards, the shuffled scar family distance 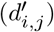 was calculated with the same way as above. The observed scar family distance was then normalized into the z-score 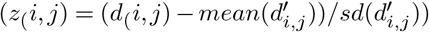. A z-score which is significantly larger than zero indicates a significantly large scar family composition difference between two clusters, implying fate commitment happened when scarring was induced.

To capture the global commitment signal, the distribution of observed z-scores across all cluster pairs was subtracted by the average distribution of the 1,000 shuffling-based results, resulting in the excess of frequency at different z-score. If significant excess of frequency is observed for positive z-score, it indicates more cluster pairs show significantly different scar family composition than random, therefore implies significant cell fate commitment happened when scars were induced. To identify cell clusters sharing similar scar family composition, a hierarchical clustering was applied to the cell clusters of interest. The input distance matrix was defined as (*D_i,j_* = *z_i,j_* − *min*(*z_i,j_*)).

### Quantification of barcode family composition similarity between clusters

The barcode family composition similarity between two cell clusters was quantified as the number of cell pairs, with each cell in one cluster, that are of the same barcode family (denoted as *n_i,j_*, Fig. 3a). To control for confounding factors including cell numbers in clusters and organoid composition, 100 random shuffling of barcode family information of cells in each organoid were applied and the random composition similarity between two clusters was estimated in the same way (denoted as 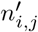). The observed barcode family composition similarity was then normalized into the z-score 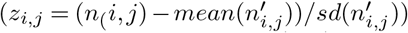. A z-transformation was further applied to scale the resulting z-scores between different cluster pairs (denoted as 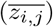), and two cutoffs (0.01 and 0.99 quantile of the standard normal distribution, i.e. *z_cutoff−_* = −2.33 and *z_cutoff_*_+_ = 2.33) were applied to get cluster pairs with significant similar 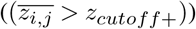 or different 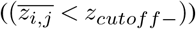 barcode family composition. To identify groups of cell clusters with similar barcode family composition, a hierarchical clustering was applied, with the input distance matrix defined as (*d_i,j_* = *max*(*z_i,j_*) −*z_i,j_*).

Alternatively, a hierarchical clustering was applied to the binomial-based normalized barcode family composition similarity distance matrix, which only takes into account the sizes of cell clusters. In brief, assuming the total number of cell pairs from the same barcode family being *N*, and two clusters *i* and *j* representing proportions of *p_i_* and *p_j_* of the whole data set, the expected number of cell pairs in these two different clusters from the same barcode family is 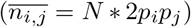 with the expected standard deviation of 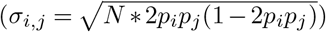, based on binomial distribution. The observed barcode family composition similarity was then normalized into the alternative z-score (*z_i,j_* = (*n_i,j_* − *n_i,j_*)/*σ_i,j_*). A hierarchical clustering was then applied to identify groups of cell clusters with similar barcode family composition, with the input distance matrix defined as (*d_i,j_* = *max*(*z_i,j_*) − *z_i,j_*) with the alternative z-scores.

### Microdissection of single organoid regions

A 200um slice of a single organoid (Org13) was cut with a vibratome. Regions that were spatially distinct were selected and microdissected away from the slice. Each microdissected area was dissociated (as described above) adjusting times and volumes to account for smaller tissue input. Following single-cell isolation, cells were captured using 10x Chromium targeting 17,000 cells per region across four separate reactions so that each region was split into two capture reactions. Transcriptome, barcode, and scar libraries were prepared as described above. All libraries were pooled and sequenced on Illumina NovaSeq S1.

The resulting sequencing reads were aligned and preprocessed as described above. Seurat (v3.1) was applied for log-normalization and highly variable genes identification (vst method, 5,000 genes, Supplementary Table 6). PCA was applied to the z-transformed expression levels of the highly variable genes, with the top-20 PCs used for UMAP embedding construction and louvain clustering (with resolution 0.6). Cell cluster annotation was done by combining canonical marker gene expression and CSS projection of cells to the whole-organoid scRNAseq data described above. In brief, the whole-organoid CSS representations of cells in the microdissected scRNAseq data were calculated as the normalized Spearman correlation between cell transcriptome and the average transcriptome of cell clusters in the whole-organoid scRNAseq data. The k-nearest neighbors (kNN, k=50) of each cell in the whole-organoid scRNAseq data were identified, as those with the shortest Euclidean distances at CSS representations. The major cell cluster label of the identified neighbors were assigned to the query cell in the microdissected scRNAseq data as the transferred label, to assist annotation of the microdissected scRNAseq data.

Barcode family composition similarities between cell clusters were quantified as described above. Hierarchical clustering was applied to the random barcode shuffling based quantification (ward.D2 method) to group cell clusters into three groups. Scar family composition differences between cell clusters in each of the three groups were then quantified as described above.

### Spatial transcriptomics

Org14 was embedded in pre-chilled Optimal Cutting Temperature (OCT). The sample was then set into a dry ice bath with isopentane until frozen and stored at −80°C. Cryosections were cut at 10um thickness, adhered to ST slides (10x) and stored at −80°C until the following day. Tissue slices were fixed in cold methanol, before being stained in Hematoxylin and eosin. ST slides were imaged as recommended on a Nikon T2i at 20x using a tile scan over all slice sections. Following image capture, tissue slices were permeabilized. Optimal permeabilization conditions were determined by using the Tissue Optimization Kit (10x) and was found to be 24 minutes. Spot-captured RNA was reverse transcribed before second strand synthesis and cDNA denaturation. qPCR was used to determine the optimal number of cDNA amplification cycles as recommended by the manufacturer. cDNA was amplified using 17-18 cycles, before continuing to Visium spatial gene expression library construction. Visium libraries were sequenced on Illumina NovaSeq SP following sequencing recommendations. Barcode and scar libraries were sequenced on NextSeq mid-output.

The resulting sequencing reads were aligned using Space Ranger for the regular Visium libraries (10x Genomics). Sections one and three were automatically tissue aligned, whereas we manually annotated tissue covering spots with Loupe Browser (10x Genomics) for section three. Spots not covering tissue were discarded manually in R (see Data/Code availability). Barcodes and scars were called using methods described above with one exception. In order to use Cell Ranger to map barcode and scar sequencing reads to the custom reference the Cell Ranger barcode whitelist was replaced with the whitelist of the Space Ranger barcode set.

Spots were annotated using CIBERSORTx^19^, where we digitally sorted each spatial spot into fractions of cell types present. To this end, we first constructed a signature matrix for deconvolution using the highly variable genes and subset of cells across all cell annotations from the whole-organoid analysis (Supplementary Table 7). We then input each detected spot across all tissue sections (S1-S3) for sorting which resulted in a matrix of spots verses each cell annotation where each row sumed to 1. We plotted the distribution of the highest proportion (score) for each spot and set a threshold so that all spots with the highest contributing proportion less than the first quartile (.405 or 40.5%) were called “unassigned”. The remaining spots were then assigned the corresponding cell annotation of their highest contributing proportion.

To quantify the relationship between barcode composition differences and spatial proximity between spots, we firstly defined barcode composition similarity between any two detected spots *i* and *j* as the Jaccard index (*J_i,j_*) of detected barcodes in the two spots, i.e. the ratio of shared barcode number to union barcode number. Barcode composition distance was then defined as (*d_i,j_* = 1 − *J_i,j_*). Correlation between the barcode composition distances and spatial distances was next calculated. Alternatively, spot pairs were grouped into two groups: 1) spot pairs with at least one barcode shared, and 2) spot pairs with no overlapping detected barcode. Spatial distances between spot pairs in the two groups were compared using two-sided Wilcoxon’s rank sum test.

### Lightsheet imaging and tracking of cerebral organoid

We generated organoids using iPSCs expressing the FUS protein tagged with EGFP, that uniformly labels the nuclei. The FUS-mEGFP (Cell Line ID: AICS-0080 cl.69) and WTC lines (Cell Line ID: GM25256) used for imaging were procured from the Coriell Institute. Organoids were imaged with the LS1 Live lightsheet microscope developed by Viventis Microscopy Sàrl (Lausanne), using a 25x objective demagnified to 18.5x and with a field of view that is 700μm. The successive z steps were acquired every 2μm for 150 steps. The frame rate for acquisition was 30 minutes and in total 100 hours of development (200 frames) were used for tracking. For imaging, EBs were embedded in a neural induction medium together with matrigel. The lightsheet data was converted into HDF5 format and visualised using the BigDataViewer^26^ in Fiji^21^. In total four nuclei were tracked, the first one for 100 hours and the next three for 65 hours. Three neighbouring nuclei were tracked in the same developing lumen area and one in another diametrically opposite location surrounding another lumen in Org15. The nuclei are continually tracked in 3D using a new large-scale tracking and track-editing framework Mastodon (preview), a next generation software of the successful tools^27,28^ developed by the Tomancak lab at the MPI-CBG (https://sites.imagej.net/Mastodonpreview/) as a plugin in Fiji. It allows semi-automated tracking and manual curation of the nuclei tracks. During lumen expansion and growth, some nuclei tracks are prematurely terminated when they move out of focus.

## Extended Data Figures

**Extended Fig. 1.**
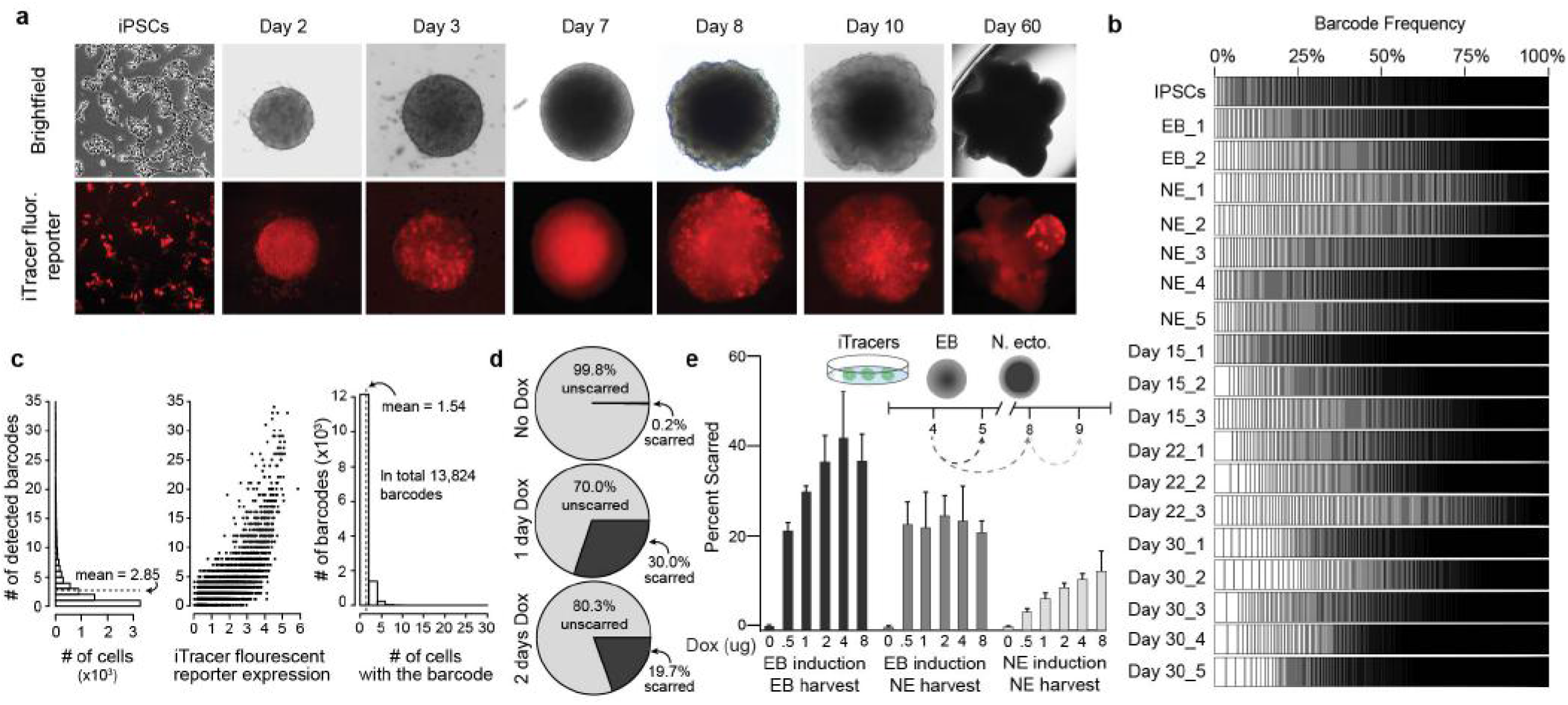
Assessment of iTracer readouts iPSC and iPSC-derived cerebral organoids. a) Brightfield and fluorescence imaging of iTracer reporter (RFP) in organoids during cerebral organoid development. b) Stacked barplots showing frequency of unique barcodes detected in bulk targeted amplicon sequencing libraries throughout cerebral organoid development. c) Barplots of the number of iTracer barcodes detected from single-cell transcriptomes. The left panel shows frequencies of cells with different numbers of detected barcodes. The dashed line shows the average number of detected barcodes per cell (2.85). The middle panel shows numbers of barcodes in relation to fluorescent reporter detection. Each dot represents one cell. The right panel shows frequencies of barcodes detected in different numbers of cells. The dashed line indicates that on average one barcode is detected in 1.54 cells. d) Scar detection in iPSCs treated with no doxycycline, 2ug of doxycycline for one day, and 2ug of doxycycline for two days. e) Stacked barplots showing frequencies of different scars detected in 3D cultures treated with 0-8ug of doxycycline at EB and Neuroectoderm stages.

**Extended Fig. 2.**
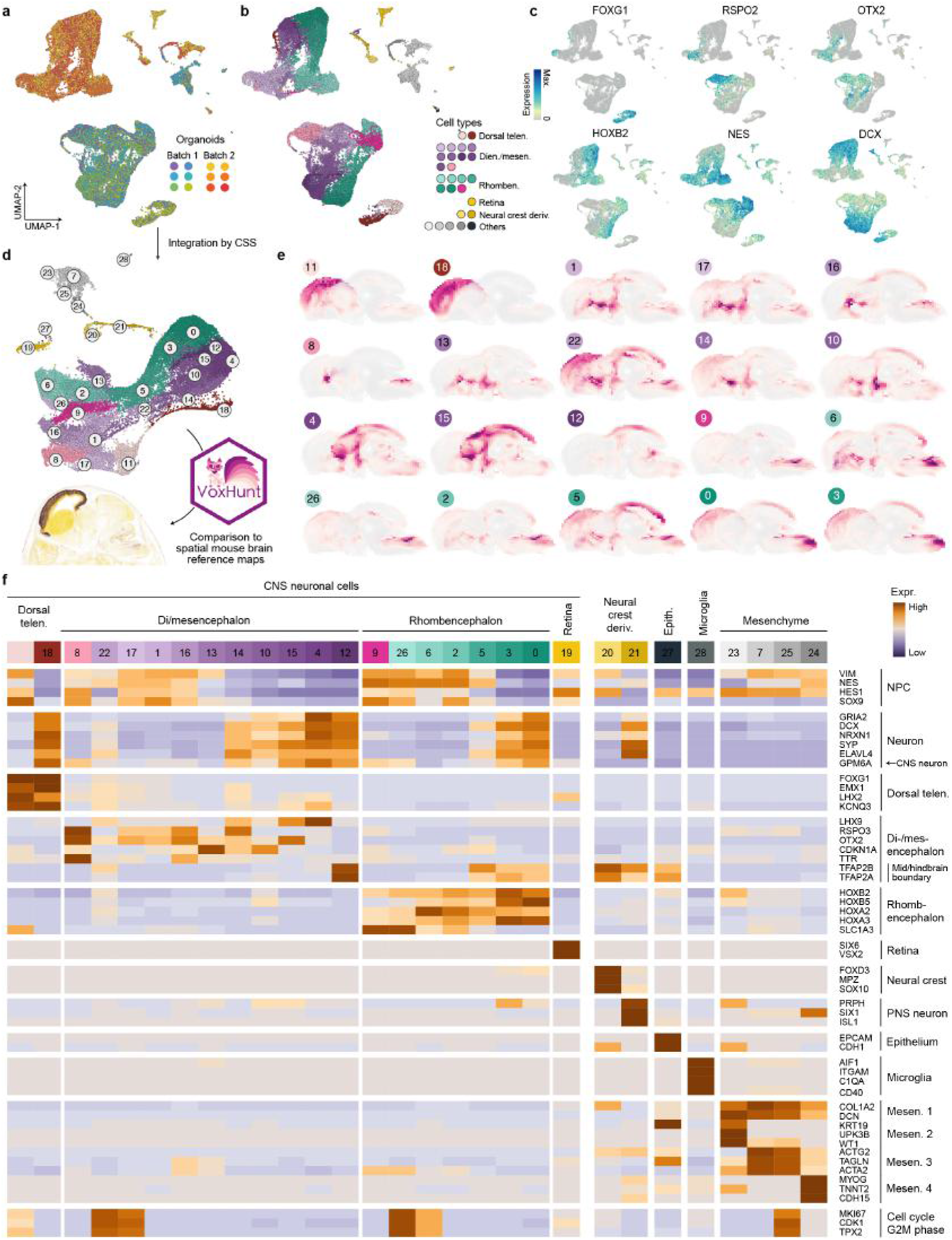
Integration and cell type annotation of iTracer cerebral organoid single-cell transcriptomes. a) UMAP embedding of single-cells from two batches of iTracer whole-organoids without integration, colored by organoid. b) UMAP embedding of single-cells from two batches of iTracer whole-organoids colored by cell type annotation (telen. – telencephalon; dien./mesen. – diencephalon/mesencephalon; rhomben. – rhombencephalon; neural crest deriv. – neural crest derivatives). c) UMAP colored by expression of selected marker genes. d) Schematic of cluster annotation of CSS integrated whole-organoid data using VoxHunt. Similarity scores are calculated between average gene expressions of identified clusters in the whole-organoid data and in situ hybridization signals in the E13.5 mouse brain in Allen Brain Atlas. e) Sagittal projections colored by scaled similarity scores of each cluster from integrated whole-organoid data to voxel maps of the E13.5 mouse brain. f) Heatmap of cell type and cell state marker genes across all CSS integrated whole-organoid clusters (mesen. – mesenchyme).

**Extended Fig. 3.**
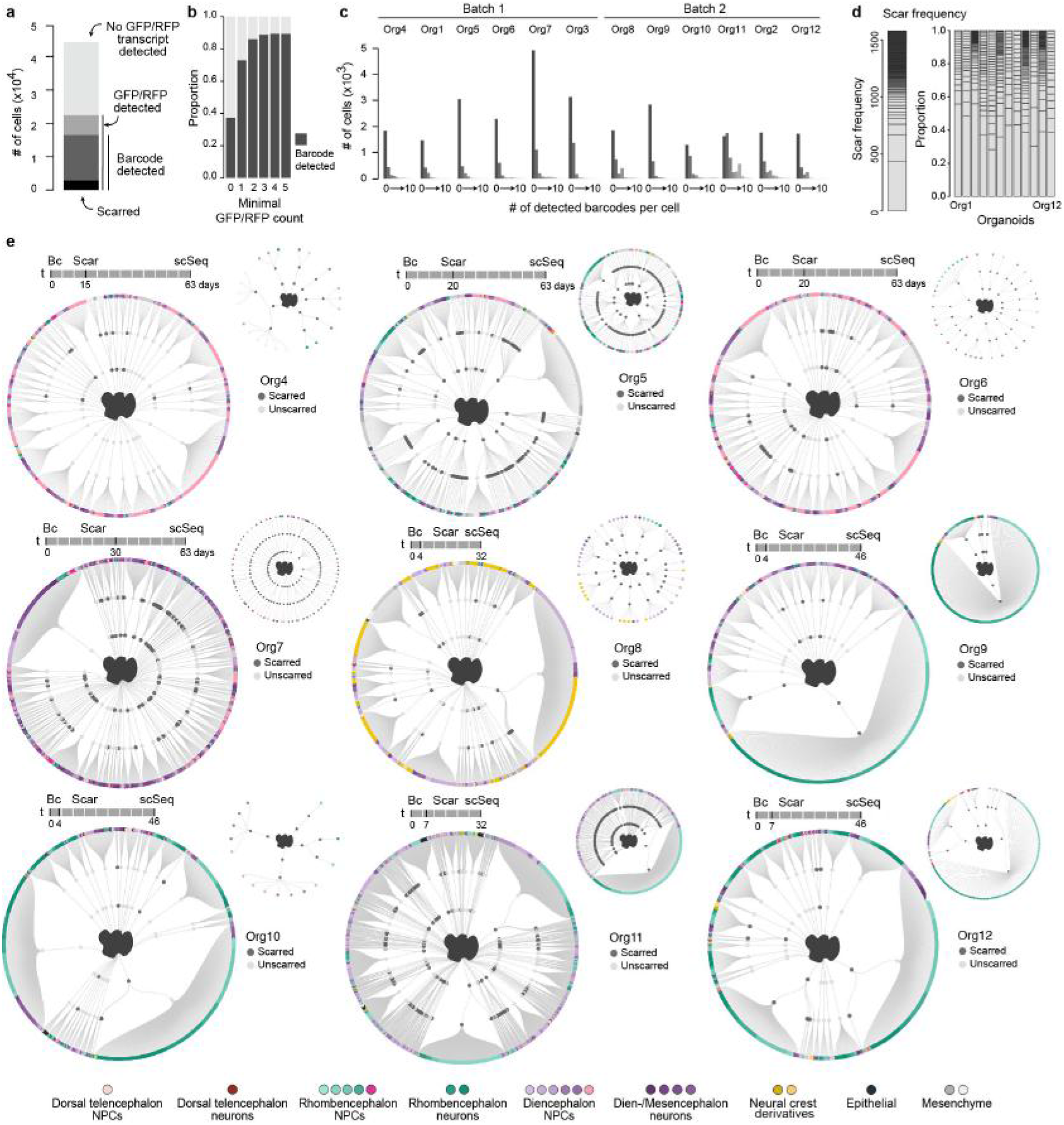
iTracer readouts enable construction of cell lineage trees from cerebral organoids. a) Stacked barplot showing the number of cells measured across all 12 organoids (lightest grey), where only iTracer reporter was captured (light grey), with reporter and barcodes (dark grey) or reporter, barcodes and scars (black). b) Stacked barplot showing the proportion of cells with barcode detected under different iTracer reporter transcript cutoffs. c) Histograms of the cell numbers with different numbers of barcodes detected across all 12 organoids. d) Stacked barplots of overall scar frequency of all scars detected across all 12 organoids (left) and their proportions in each organoid separately (right). e) Lineage plots show full lineage reconstructions, as well as the subset of cells where scars were detected from 9 organoids. The first and second order deviation nodes represent barcode and scar families respectively, with the terminal branches indicating individual cells. Each cell is colored based on the cell type designation.

**Extended Fig. 4.**
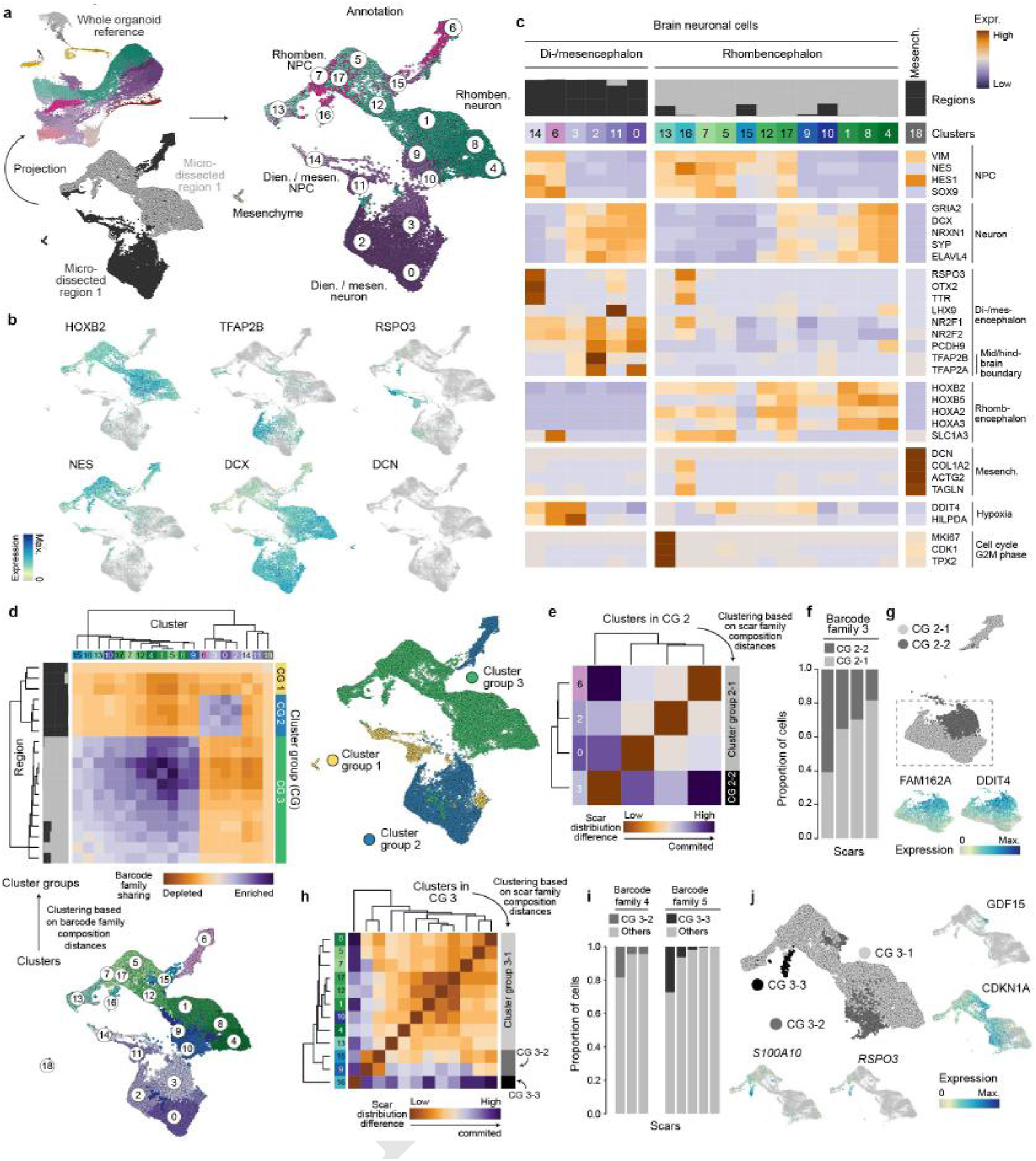
Deep cell and lineage sampling of micro-dissected regions from cerebral organoids. a) Schematic of data projection procedure for single-cells from two regions of a micro-dissected organoid to the whole-organoid scRNAseq data to assist cell annotation. UMAP embedding of single cells colored by projected annotation from the whole-organoid scRNAseq data. b) Expression of selected cell type markers in cells in the two regions of the micro-dissected organoid. c) Heatmap of cell type and cell state marker genes across all micro-dissected organoid clusters. d) Hierarchical clustering of the micro-dissected organoid clusters, based on the barcode family composition distances. UMAP embedding of single-cells is colored by the resulting three groups of clusters (CG.1-CG.3). Clusters in each group share similar barcode family compositions. e) Hierarchical clustering of cluster group 2 (CG.2) based on scar family composition distance, to identify two subgroups of clusters. Clusters in each of the subgroups share similar scar family compositions. f) Stacked barplots showing distributions of cell proportions across subgroups of CG.2 clusters with distinct scar family compositions. Each stacked bar shows a different scar family in the same example barcode family 3. g) UMAP embedding of cells in CG.2 clusters, colored by the two cluster subgroups with distinct scar family compositions, and expression of example genes with differential expression between the two subgroups. h) Hierarchical clustering of cluster group 3 (CG.3) based on scar family composition distance, to identify three subgroups of clusters. Clusters in each of the subgroups share similar scar family compositions. i) Stacked barplots showing distributions of cell proportions across subgroups of CG.3 clusters with distinct scar family compositions. Each stacked bar shows a different scar family in the same example barcode family 4 (left) or barcode family 5 (right). j) UMAP embedding of cells in CG.3 clusters, colored by the three cluster subgroups with distinct scar family compositions, and expression of example genes with differential expression between the two subgroups.

**Extended Fig. 5.**
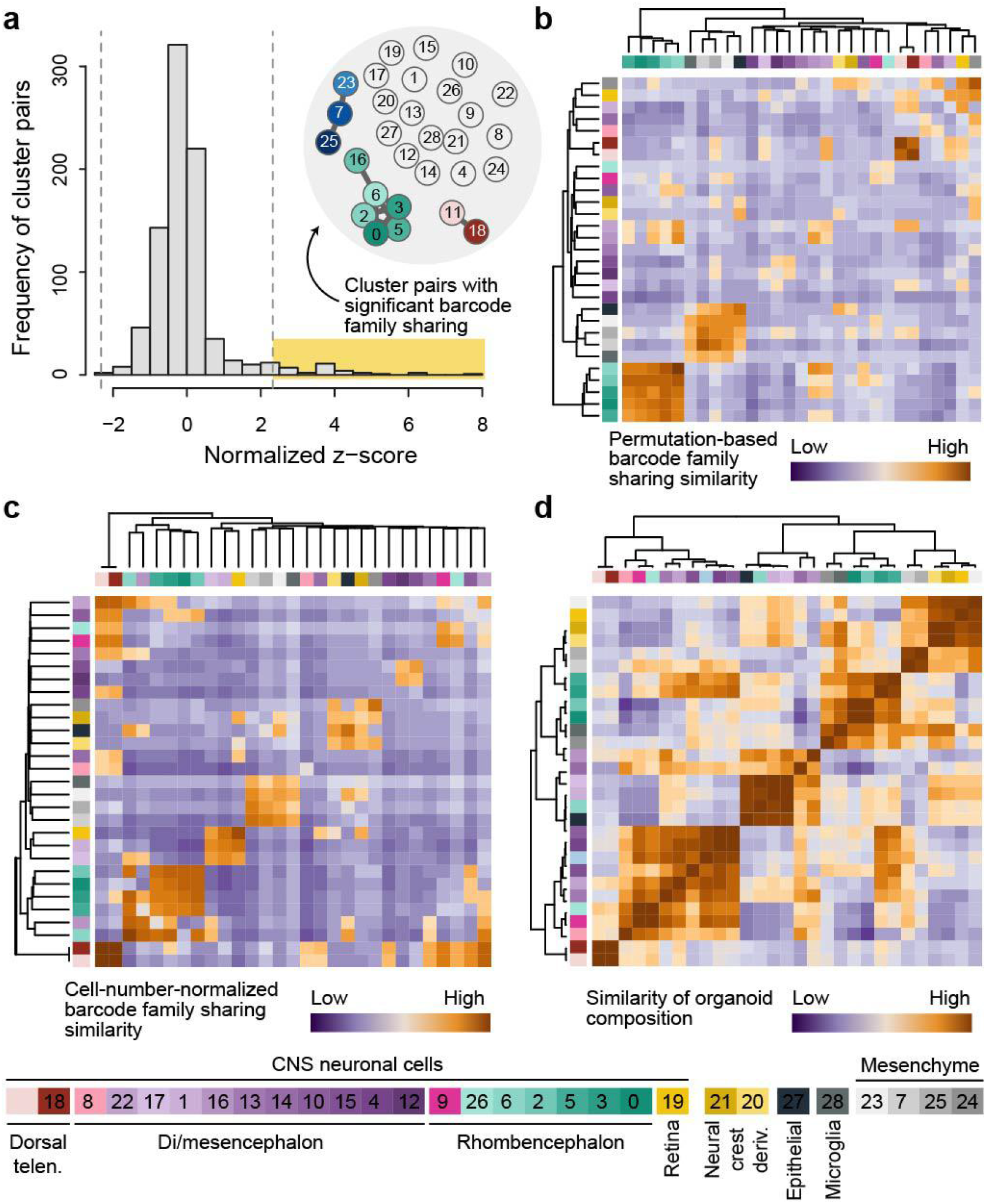
iTracer barcode families accumulate in distinct brain regions. a) Histogram of the distribution of all normalized z-scores. Cutoffs are the 99% quantile of the standard normal distribution (2.32), and 1% quantile (−2.32). b) Heatmap of permutation based barcode family similarity, with hierarchical clustering (ward.D2 method) applied to permutation-based barcode family composition distances between clusters. c) Heatmap of cell number normalized barcode family similarity, with hierarchical clustering (ward.D2 method) applied to cell-number-normalized barcode family composition distances. d) Heatmap of similarity of organoid composition, defined as the Pearson’s correlation coefficient between cell frequencies of each cluster across different organoids, with hierarchical clustering (ward.D2 method) applied to the correlation distances between cell frequencies of clusters across different organoids.

**Extended Fig. 6.**
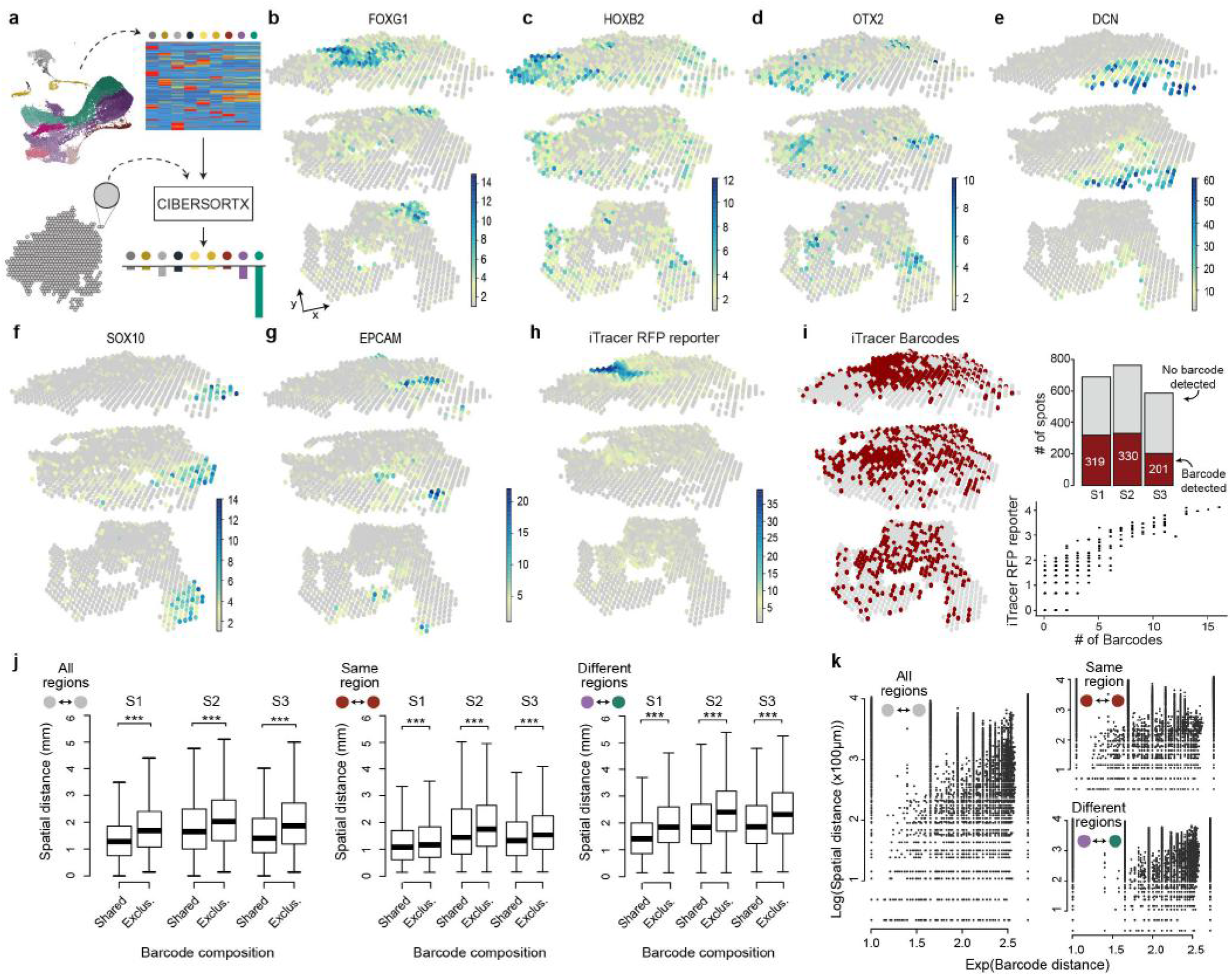
Regionalization of cell types and iTracer barcodes across cerebral organoids. a) Schematic of spot annotation by digital cytometry using CIBERSORTx. b-g) 3D spatial feature plots of expression of genes marking different cell types and brain regions. h) 3D spatial feature plots of expression of iTracer RFP reporter. i) 3D spatial plot of spots with (red) and without (grey) iTracer barcodes detected. Barplot of detected iTracer barcodes across all sections. Scatterplot of detected barcodes and iTracer RFP reporter. j) Boxplots of the iTracer barcode composition of each spot pair vs the spatial distance of each spot pair across all sections (S1-S3) in any regions (left), same cell type regions (middle) and different cell type regions (right). Two-sided Wilcoxon rank sum tests were performed comparing shared to same and exclusive groups, *** indicates p-values<0.0001. k) Scatter plots of barcode distance between each spot pair at the same section vs their spatial distance, in all the three sections, in any regions (left), same cell type regions (top-right) and different cell type regions (bottom-right).

**Extended Fig. 7.**
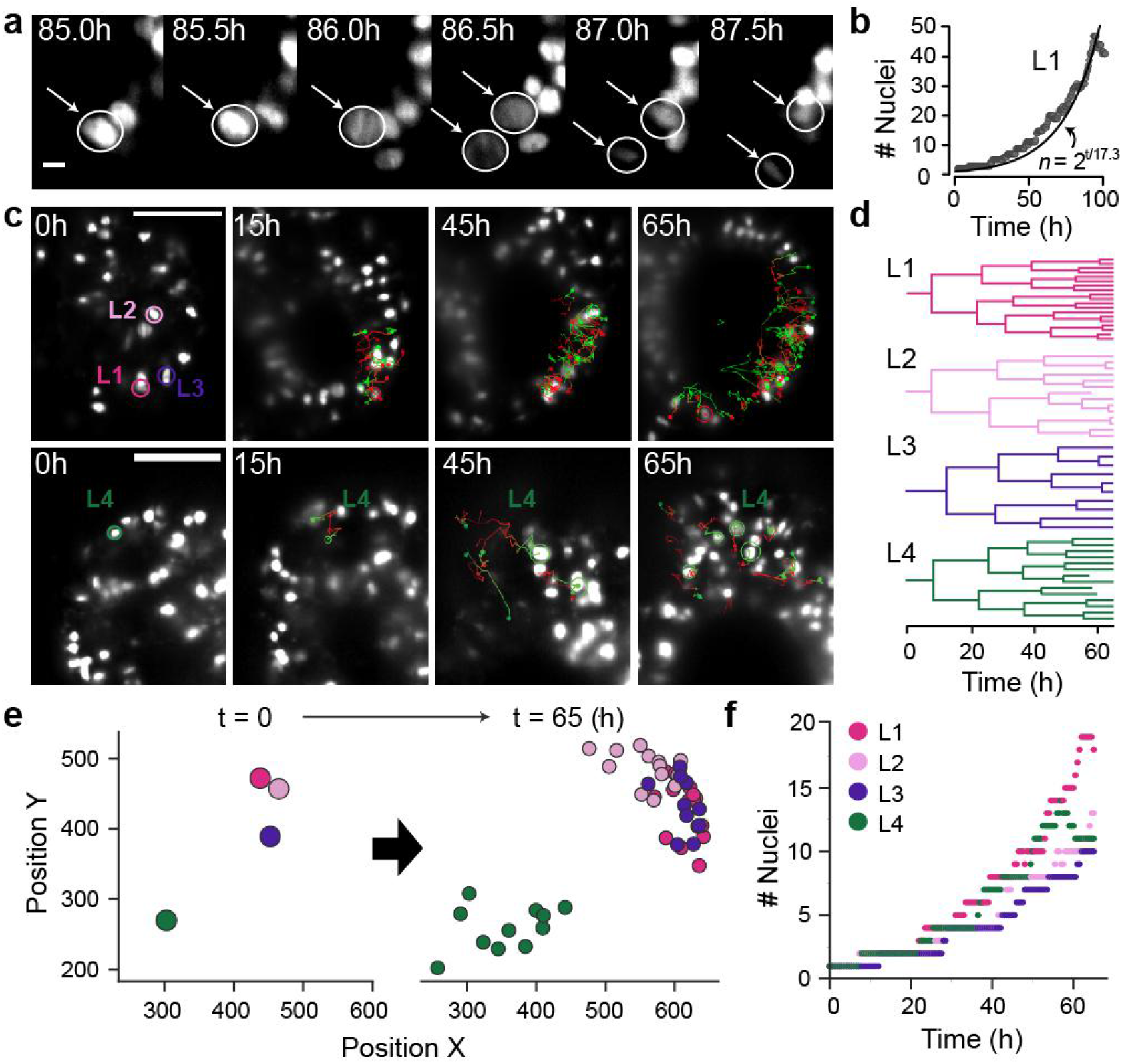
Tracking spatial distribution of nuclei lineages using Light sheet microscopy. a) Example images show a nucleus dividing into two daughter nuclei in the developing organoid. White circles represent the manual detection and tracking over time. b) Scatter plot shows increase in the number of nuclei over 100 hours of tracking in lineage one (L1). The curve shows the exponential model estimated by the data, with the estimated doubling time being 17.3 hours. c) Nuclei tracking of three lineages (L1-L3) in the same lumen area (top) and of a fourth lineage (L4) surrounding a diametrically opposite lumen (bottom). d) Lineage trees for the four tracked nuclei lineages. e) Spatial distribution in the x-y plane of the four tracked lineages. Nuclei are shown at 0 hours and 65 hours, colored by lineage. f) Dotplot shows the increase in number of daughter nuclei for all four lineages over 65 hours.

