## Supplementary Materials for "Lineage recording reveals dynamics of cerebral organoid regionalization"

### SUPPLEMENTARY TABLES

**Supplementary Table 1: Overview of organoids used in this study.**

**Supplementary Table 2: Metadata of cells in the scRNAseq data of the whole cerebral organoids.**

**Supplementary Table 3: Metadata of cells in the scRNAseq data of the microdissected cerebral organoid regions.**

**Supplementary Table 4: Metadata of spots in the Spatial Transcriptomics (10X Genomics Visium) data of cerebral organoid sections.**

**Supplementary Table 5: Oligo and primer sequences used in this study.**

**Supplementary Table 6: Highly variable genes used in this study.** Highly variable genes were determined for each data set separately using Seurat (v3.1, vst method).

**Supplementary Table 7: Signature matrix used by CIBERSORTx to deconvolve the transcriptomes of visium spots.** The genes were selected by CIBERSORTx from the highly variable genes identified in the scRNAseq data of whole cerebral organoids.

### SUPPLEMENTARY MOVIES

**Supplementary Movie 1: Cross section (x-y) from a time lapse recording of a developing cerebral organoid imaged for 100 hours using light-sheet microscopy.** The frame rate for acquisition was 30 minutes. Labeled nuclei (FUS-mEGFP) are colored in green. Scale bar is 100 $\mu$m. Time-stamp is hh:mm.

**Supplementary Movie 2: Time lapse video showing a cross section of a lumen of the organoid in supplementary movie 1.** Arrows mark a nucleus dividing into two daughter nuclei. Time-stamp is hh:mm and 00:00 corresponds to the beginning of the supplementary movie 2, which is 83 hours of organoid development. Scale bar is 100 $\mu$m.

**Supplementary Movie 3: Movie showing the 3D position of tracked nuclei in lineage one (L1) over the complete 100 hours time course.** Every black dot represents a daughter nucleus in the lineage at one time point. The pink dots highlight all nuclei in one selected branch of the tree.

**Supplementary Movie 4: Movie showing the 3D position of tracked nuclei for all four lineages over a time course of 65 hours.** The nuclei belonging to each lineage (L1-L4) are distinctly colored with the green lineage originating in a different lumen area compared to the other three lineages.
